# Life histories and niche dynamics in late Quaternary proboscideans from Midwestern North America: evidence from stable isotope analyses

**DOI:** 10.1101/2020.01.08.896647

**Authors:** Chris Widga, Greg Hodgins, Kayla Kolis, Stacey Lengyel, Jeff Saunders, J. Douglas Walker, Alan D. Wanamaker

## Abstract

Stable isotopes of mammoths and mastodons have the potential to illuminate ecological changes in late Pleistocene landscapes and megafaunal populations as these species approached extinction. The ecological factors at play in this extinction remain unresolved, but isotopes of bone collagen (δ^13^C, δ^15^N) and tooth enamel (δ^13^C, δ^18^O, ^87^Sr/^86^Sr) from the Midwest, USA are leveraged to examine ecological and behavioral changes that occurred during the last interglacial-glacial cycle. Both species had significant C3 contributions to their diets and experienced increasing levels of niche overlap as they approached extinction. A subset of mastodons after the last glacial maximum (LGM) exhibit low δ^15^N values that may represent expansion into a novel ecological niche, perhaps densely occupied by other herbivores. Stable isotopes from serial and micro-sampled enamel show increasing seasonality and decreasing temperatures as mammoths transitioned from Marine Isotope Stage (MIS) 5e to glacial conditions (MIS 4, MIS 3, MIS 2). Isotopic variability in enamel suggests mobility patterns and life histories have potentially large impacts on the interpretation of their stable isotope ecology. This study further refines the ecology of midwestern mammoths and mastodons demonstrating increasing seasonality and niche overlap as they responded to landscape changes in the final millennia before extinction.

## INTRODUCTION

Historically, the late Quaternary record of mammoths and mastodons in the Midwest has played an important role in understanding megafaunal extinctions in North America (e.g., Fisher 2008, 2018; Graham et al., 1981; Saunders et al. 2010; Widga et al., 2017a; Yansa and Adams, 2012). Whether extinctions are viewed as human-induced (Mosimann and Martin 1975; Alroy 2001; Surovell et al. 2005; Fisher 2009; Surovell et al. 2016), the result of late Pleistocene landscape changes (Stuart et al. 2004; Nogués-Bravo et al. 2008; Widga et al. 2017a), a function of other ecological processes (Ripple and Van Valkenburgh 2010), or some combination, this region is unparalleled in its density of large, late Quaternary vertebrates and associated paleoecological localities. For these reasons, the midcontinent has proven critical to understanding ecological dynamics in proboscidean taxa leading up to extinction. In recent years, it has been recognized that multiple proboscidean taxa shared an ecological niche in late Pleistocene landscapes (Saunders et al. 2010) despite distinct extinction trajectories (Widga et al. 2017a). Chronological studies indicate that proboscideans in this region experienced extinction *in situ,* rather than mobilizing to follow preferred niche space (Saunders et al. 2010). Despite refinements in our understanding of mammoths and mastodons in the region, many significant questions remain. Modern megafauna have a profound impact on local vegetation (Guldemond and Van Aarde 2008; Valeix et al. 2011), and it is unclear what effect late Pleistocene mammoths and mastodons would have had on canopy cover, nutrient cycling, and fruit dispersal in non-analogue vegetation communities. There are still issues of equifinality in how the presence of human predators (Fisher 2009) or the virtual absence of both humans and large carnivores (Widga et al. 2017a) impacted proboscidean populations in the region as they neared extinction.

The high profile debate surrounding the cause of late Pleistocene megafaunal extinctions has spurred a number of productive regional studies to address the timing and paleoecology of extinction in megafaunal taxa (Pacher and Stuart 2009; Stuart and Lister 2011, 2012; Stuart 2015; Widga et al. 2017a). The results of this research serve to constrain the number of possible extinction scenarios and to highlight the need for regional-scale and taxon-specific analyses.

As in modern ecosystems, Proboscidea in late Pleistocene North America were long-lived taxa that likely had a profound impact on the landscape around them. Due to their size and energetic requirements, elephantoids are a disruptive ecological force, promoting open canopies in forests through tree destruction (Chafota and Owen-Smith 2009) and trampling vegetation (Plumptre 1994). Their dung is a key component of soil nutrient cycling (Owen-Smith 1992; Augustine et al. 2003), and was likely even more important in N limited tundra and boreal forest systems of temperate areas during the late Pleistocene. Even in death, mammoths and mastodons probably wrought major changes on the systems within which they were interred (Coe 1978; Keenan et al. 2018) providing valuable nutrient resources to scavengers and microbial communities. Precisely because proboscideans play such varied and important roles in the ecosystems they inhabit, they are good study taxa to better understand Pleistocene ecosystems. They are also central players in many extinction scenarios.

One of the major challenges to ecological questions such as these is scale (Delcourt and Delcourt 1991; Denny et al. 2004; Davis and Pineda-Munoz 2016). Processes that are acting at the level of an individual or a locality can vary significantly in space and time (Table 1). Paleoecological data collected from individual animals from local sites is constrained by larger regional patterns that may or may not be apparent (e.g., taphonomic contexts, predator-prey dynamics). Paleoecological studies of vertebrate taxa often begin with the premise that the individual is an archive of environmental phenomena experienced in life. In long-lived taxa such as proboscideans, this window into past landscapes may span many decades. This longevity is both a benefit and a challenge to studies of proboscidean paleoecology.

**Table 1.** Scales of paleoecological analysis.

| Scale | Method | Phenomena | Selected References |
| --- | --- | --- | --- |
| Small<br>(Days) | Stomach contents, enamel micro-wear | Animal health, Landscape (local scale) | Rhodes et al. 1998; Teale and Miller 2012; Green, DeSantis, and Smith 2017; Smith and DeSantis 2018. |
| Meso<br>(Weeks-Months) | Hair, Tusk/Tooth dentin, Tooth enamel | Seasonal diets, Reproductive events (e.g., musth, calving), Migration/Dispersal | Fox and Fisher 2001; Hoppe 2004; Hoppe and Koch 2007; Cerling et al. 2009; Metcalfe and Longstaffe 2012; Fisher 2018 |
| Large<br>(Years - Decades) | Bone collagen, tooth enamel, tooth mesowear | Climate/Vegetation trends, Large-scale land-use trends, Population-level trends in diet responses to landscape changes. | Bocherens et al. 1996; Iacumin et al. 2010; Szpak et al. 2010 |

Some approaches offer relatively high-resolution snapshots of animal ecology at a scale that is of a short duration (days). Micro-wear analyses of dentin and enamel (Green et al. 2017; Smith and DeSantis 2018) are increasingly sophisticated, and have the potential to track shortterm dietary trends. The remains of stomach contents also provide ecological information at this scale (Lepper et al. 1991; Newsom and Mihlbachler 2006; van Geel et al. 2011; Fisher et al. 2012; Teale and Miller 2012; Birks et al. 2019), which are essentially the ‘last meal’ representing a few hours of individual browsing. These techniques offer paleoecological insights that are minimally time-averaged, and at a timescale that may be comparable to modern observations of animal behavior.

Other approaches resolve time periods that are weeks to months in duration. Fisher’s (Fisher and Fox 2006; Fisher 2009; 2018) work on incremental growth structures in proboscidean tusk and molar dentin reliably record weekly to monthly behaviors. The resolution of these methods may even include short-term, often periodic, life history events, such as reproductive competition (musth) and calving. Other researchers (Hoppe et al. 1999; Metcalfe and Longstaffe 2012; 2014; Pérez-Crespo et al. 2016) have explored incremental growth trends in proboscidean tooth enamel. Adult molars form over the course of 10-12 years with an enamel extension rate of ~1 cm/year in both modern elephants (Uno et al. 2013) and mammoths (Dirks et al. 2012; Metcalfe and Longstaffe 2012) providing the opportunity to understand meso-scale (potentially monthly) changes in diet and behavior.

Finally, some techniques measure animal diet and behavior over much longer scales (years-decades). In humans, bone collagen is replaced at a rate of 1.5-4% per year (Hedges et al. 2007). For equally long-lived proboscidean taxa, this means that stable isotope analyses of bone collagen is essentially sampling a moving average of ~20 years of animal growth. For younger age groups, this average will be weighted towards time periods of accelerated maturation (adolescence), when collagen is replaced at a much greater rate (Hedges et al. 2007). Tooth enamel can also be sampled at a resolution (i.e., “Bulk” enamel) that averages a year (or more) of growth (Hoppe 2004; Baumann and Crowley 2015), and it is likely that this is the approximate temporal scale that is controlling tooth mesowear (Fortelius and Solounias 2000).

Stable isotope studies are an important part of the paleoecological toolkit for understanding Quaternary proboscideans and are capable of resolving animal behavior at multiple timescales. Progressively larger, more complete datasets characterize isotopic studies of Beringian mammoths, where stable carbon and oxygen isotopes in bone collagen and tooth enamel reliably track climate and landscape changes over the late Pleistocene (Bocherens et al. 1996; Iacumin et al. 2010; Szpak et al. 2010; Arppe et al. 2019), the place of mammoths in regional food webs (Fox-Dobbs et al. 2007, 2008), and characteristics of animal growth and maturation (Metcalfe and Longstaffe 2012; Rountrey et al. 2012; El Adli et al. 2017).

These studies have also been important to understanding both lineages of proboscideans in temperate North America. From the West Coast (Coltrain et al. 2004; El Adli et al. 2015) to the southwestern (Metcalfe et al. 2011), and eastern US (Koch et al. 1998; Hoppe and Koch 2007), isotopic approaches have been very successful in understanding local to regional scale behavior in mammoths and mastodons. Although there have been efforts to understand mammoth and mastodon behaviors in the Midwest at relatively limited geographic scales (Saunders et al. 2010; Baumann and Crowley 2015), there is a need to systematically address long-term isotopic trends throughout the region. In this paper, we approach this problem from a broad regional perspective, leveraging a recently reported radiocarbon (^14^C) dataset with associated isotopic data on bone collagen. We also utilize an enamel dataset consisting of both serial bulk and micro-milled mammoth molar enamel samples (C, O, Sr isotope systems). Together, the results of these analyses offer a picture of mammoth and mastodon diets (δ^13^C, δ^15^N), late Quaternary paleoclimate (δ^18^O), and animal mobility (^87^Sr/^86^Sr) that is geographically comprehensive and spans the past 50,000 years.

## MATERIALS AND METHODS

The proboscidean dataset in this study (Figure 1; SM Table 1, SM Table 2, SM Table 3) was acquired with the goal of understanding mammoth and mastodon population dynamics during the late Pleistocene as these taxa approach extinction. Chronological and broad-scale paleoecological implications for this dataset were explored in Widga et al. (2017a) and more recently in Broughton and Weiztel (2018). In this paper we focus on the implications of these data for the stable isotope ecology of midwestern Proboscidea. We also discuss annual patterns in five serially sampled mammoth teeth spanning the last glacial-interglacial cycle. Finally, we micro-sampled two mammoths from the Jones Spring locality in Hickory Co., MO. Although beyond the range of radiocarbon dating, both samples are associated with well-dated stratigraphic contexts (Haynes 1985). Specimen 305JS77 is an enamel ridge-plate recovered from unit d1 (spring feeder), refitting to an M3 from unit c2 (lower peat). Unit c2 is part of the lower Trolinger formation (Haynes 1985) and can be assigned to Marine Isotope Stage (MIS) 4. Specimen 64JS73 is an enamel ridge-plate from unit e2 (sandy peat) in the upper Trolinger formation (Haynes 1985) and can be assigned to MIS 3. Together, these samples provide dozens of seasonally-calibrated isotopic snapshots representing mammoth behavior from individuals that pre-date the last glacial maximum (LGM).

**Figure 1.**
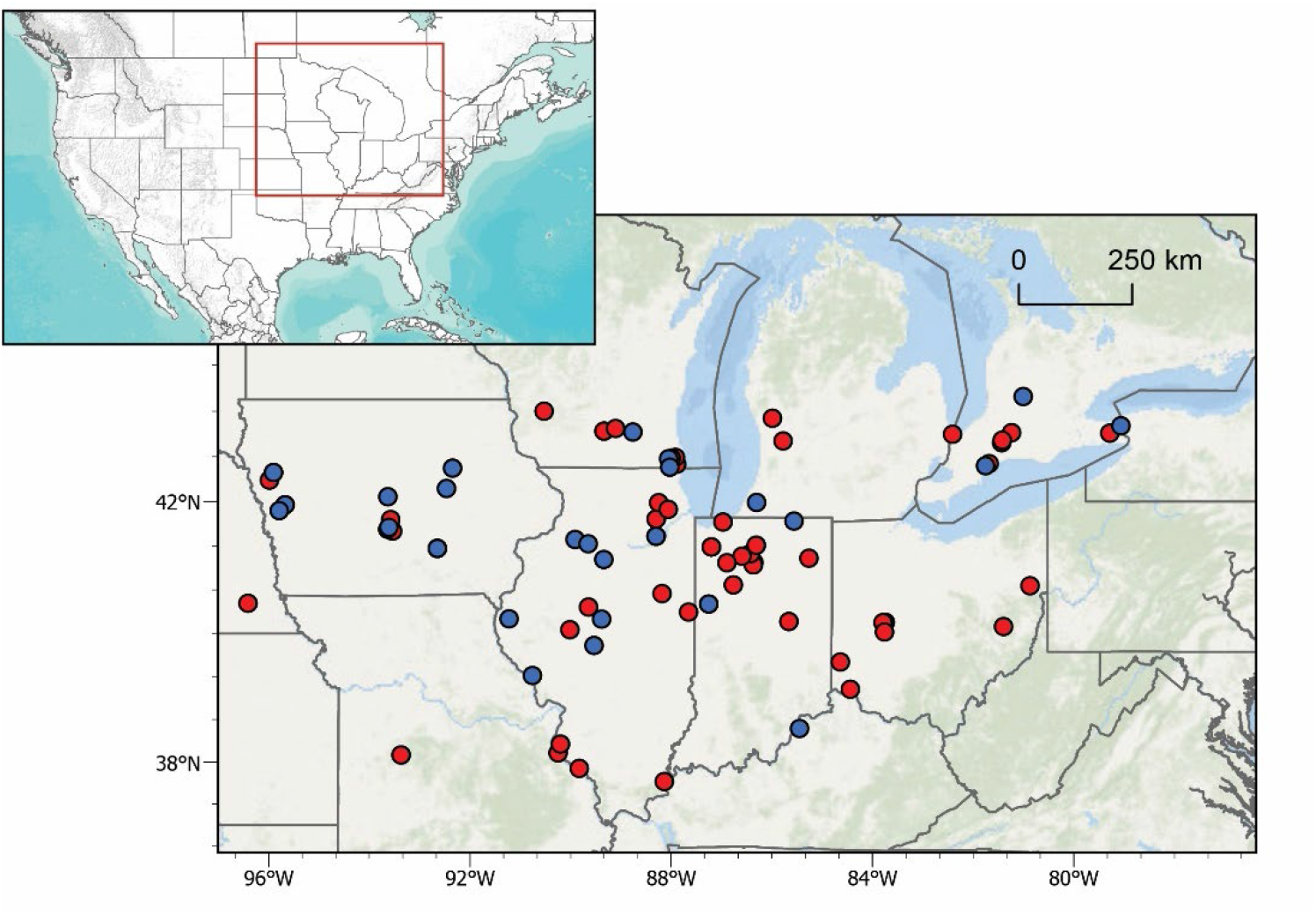
Map of dated midwestern mammoths (blue) and mastodons (red) with associated δ^13^C_coll_ and δ^15^N_coll_ data. See SM Table 2 for details.

**Table 2.** Mastodon and Mammoth collagen stable isotope values, by chronozone. δ^13^C values reported as ‰ relative to VPDB. δ^15^N values reported as ‰ relative to Air.

| Taxon | Chronozone | N | $\bar{x} \delta^{13}\text{C}$ (s.d) | $\bar{x} \delta^{15}\text{N}$ (s.d) |
| --- | --- | --- | --- | --- |
| *Mammut* | Younger Dryas (12.9-11.5 ka) | 2 | -20.6 (0.4) | 2.9 (0.6) |
|  | Allerød (14.0-12.9 ka) | 35 | -21.1 (0.6) | 4.2 (2.1) |
|  | Bølling (14.6-14.0 ka) | 6 | -20.4 (0.5) | 3.8 (1.5) |
|  | Oldest Dryas (14.6-17.4 ka) | 5 | -20.5 (0.7) | 4.6 (1.6) |
|  | LGM II (17.4-20.9 ka) | 0 |  |  |
|  | LGM I (20.9-22.9 ka) | 0 |  |  |
|  | Pre-LGM (>22.9 ka) | 6 | -20.9 (0.7) | 6.1 (1.3) |
|  | Average | 54 | -21.0 (0.7) | 4.4 (2.0) |
| *Mammuthus* | Allerød (14.0-12.9 ka) | 3 | -20.6 (0.9) | 6.1 (2.0) |
|  | Bølling (14.6-14.0 ka) | 1 | -19.9 | 5.8 |
|  | Oldest Dryas (14.6-17.4 ka) | 4 | -21.4 (0.5) | 7.9 (1.3) |
|  | LGM II (17.4-20.9 ka) | 4 | -21.1 (0.4) | 7.4 (1.3) |
|  | LGM I (20.9-22.9 ka) | 5 | -20.1 (1.1) | 7.6 (1.2) |
|  | Pre-LGM (>22.9 ka) | 5 | -19.6 (0.8) | 8.2 (0.7) |
|  | Average | 22 | -20.4 (1.0) | 7.5 (1.3) |

\* $\bar{x}$ = mean; s.d. = standard deviation

**Table 3.** Summary of serial and micro-sampled *Mammuthus* tooth enamel: δ^13^C, δ^18^O, and ^87^Sr/^86^Sr.

| ID | $\delta^{13}\text{C}$ | | | | $\delta^{18}\text{O}$ | | | | $^{87}\text{Sr}/^{86}\text{Sr}$ | | | |
| --- | --- | --- | --- | --- | --- | --- | --- | --- | --- | --- | --- | --- |
| Serial Analyses | X | Min | Max | Amp | X | Min | Max | Amp | X | Min | Max | Amp |
| Principia College (MIS 2) | -10.9 | -11.3 | -10.0 | 1.4 | -7.9 | -8.7 | -6.7 | 2.0 |  |  |  |  |
| Brookings Mammoth (MIS 2) | -9.9 | -10.5 | -9.4 | 1.1 | -9.2 | -10.0 | -7.8 | 2.2 |  |  |  |  |
| Schaefer Mammoth (MIS 2) | -9.0 | -9.9 | -7.0 | 3.0 | -8.3 | -9.9 | -6.6 | 3.3 |  |  |  |  |
| Jones Sp. Mammoth, 243JS75 (MIS 4) | -1.7 | -2.3 | -0.2 | 2.1 | -3.4 | -4.5 | -1.6 | 2.9 |  |  |  |  |
| Jones Sp. Mammoth, 232JS77 (MIS 5e) | -2.8 | -3.8 | -1.3 | 2.5 | -0.5 | -4.6 | 3.6 | 1.1 | 0.7162 | 0.7146 | 0.7177 | -0.0031 |
| Micro-Analyses |  |  |  |  |  |  |  |  |  |  |  |  |
| 305JS77 (MIS 5e) | -2.4 | -3.5 | -0.2 | 3.3 | -3.9 | -4.9 | -2.9 | 2.0 | 0.7109 | 0.7106 | 0.7118 | -0.0012 |
| 64JS73 (MIS 3) | -2.8 | -4.3 | -1.7 | 2.6 | -4.9 | -12.0 | -0.4 | 11.7 | 0.7152 | 0.7140 | 0.7164 | -0.0025 |

### Mammoth and Mastodon bone collagen

Proboscidean samples were selected to widely sample midwestern Proboscidea, both stratigraphically and geographically (Widga et al. 2017a). Due to extensive late Pleistocene glaciation in the region, this dataset is dominated by samples dating to the LGM or younger (<22 ka). Only 14 out of 93 (15%) localities predate the LGM.

All samples were removed from dense bone, tooth or tusk dentin and submitted to the University of Arizona AMS laboratory. Collagen was prepared using standard acid-base-acid techniques (Brock et al. 2010), its quality evaluated visually, and through ancillary Carbon:Nitrogen (C:N) analyses. Visually, well-preserved collagen had a white, fluffy appearance and C:N ratios within the range of modern bones (2.9-3.6) (Tuross et al. 1988). Samples outside of this range were not included in the study. Samples that had the potential to be terminal ages were subjected to additional analyses where the ABA-extracted gelatin was ultrafiltered (UF) through >30 kD syringe filters to isolate relatively undegraded protein chains (Higham et al. 2006). This fraction was also dated. All radiocarbon ages in this dataset are on collagen from proboscidean bone or tooth dentin and available in Widga et al. (2017a), through the Neotoma Paleoecology Database (www.neotomadb.org) or in SM Table 1. Measured radiocarbon ages were calibrated in Oxcal v4.3 (Bronk Ramsey 2009) using the Intcal13 dataset (Reimer et al. 2013). All stable isotope samples were analyzed on a Finnigan Delta PlusXL continuous-flow gas-ratio mass spectrometer coupled to a Costech elemental analyzer at the University of Arizona. Standardization is based on acetanilide for elemental concentration, NBS-22 and USGS-24 for δ^13^C, and IAEA-N-1 and IAEA-N-2 for δ^15^N. Isotopic corrections were done using a regression method based on two isotopic standards. The long-term analytical precision (at 1σ) is better than ± 0.1‰ for δ^13^C and ± 0.2‰ for δ^15^N. All δ^13^C results are reported relative to Vienna Pee Dee Belemnite (VPDB) and all δ^15^N results are reported relative to N-Air.

### Serial and Micro-sampling of Mammoth tooth enamel

Mammoth enamel ridge-plates were sampled at two different scales. Serial sampling consisted of milling a series of 5-10 mg samples of enamel powder with a handheld rotary tool equipped with a 1.5 mm diameter carbide bit along the axis of growth. Sample spacing was ~1 sample per centimeter of tooth growth. However, given the geometry and timing of enamel maturation (Dirks et al. 2012), these samples at best, approximate an annual scale of dietary and water inputs.

Micro-sampling however, has the potential to record sub-annual patterns in animal movement and behavior (Metcalfe and Longstaffe 2012). For this project, we built a custom micromill capable of *in situ,* micron-resolution, vertical sampling of a complete mammoth molar. This micromill setup consisted of two Newmark linear stages coupled to a Newmark vertical stage to allow movement in 3-dimensions. These stages were controlled by a Newmark NSC-G 3-axis motion controller using GalilTools on a PC. A 4-cm diameter ball joint allowed levelling of a metal (Version 1) or acrylic (Version 2) plate for holding a specimen. The armature for Version 1 consisted of a 1971 Olympus Vanox microscope retrofitted with a stationary Proxxon 50/E rotary tool using a 0.5mm end mill. Version 2 has replaced this setup with a U-strut armature using a 3D printed drill mount to allow for greater vertical and horizontal movement to accommodate large, organically-shaped specimens. Specimens were stabilized on the mounting plate using heat-flexible thermoplastic cradle affixed to a metal plate with machine screws (Version 1). However, we later developed an acrylic mounting plate method where a mammoth tooth could be sufficiently stabilized using zip ties (Version 2). This micromill was developed to address the challenges of accurately micro-milling large specimens with minimal instrumentation costs. Complete plans for this micromill are available under an open hardware license at https://osf.io/8uhqd/?view_only=43b4242623a94a529e4c6ef2396345e9.

Micro-mill sampling resolution was 1 sample per millimeter along the growth axis of the tooth plate (Figure 2). Each sample was milled in 100μm-deep passes through the entire thickness of the enamel. The lowest enamel sample (i.e., closest to the enamel-dentin boundary) in the series was used for isotopic analyses to minimize the effects of mineralization and diagenesis on the biological signal (Zazzo et al. 2006). Enamel powder was collected in deionized water to; 1) maximize sample recovery, and 2) lubricate the mill. These samples were too small for standard pretreatment of tooth enamel CO_3_ (Koch et al. 1997). However, paired bulk enamel samples treated with 0.1 N acetic acid and 2.5% NaOCl show results that are the same as untreated bulk samples. Although this technique is both time- and labor-intensive, it is minimally invasive and is capable of sampling enamel growth structures at high resolution.

**Figure 2.**
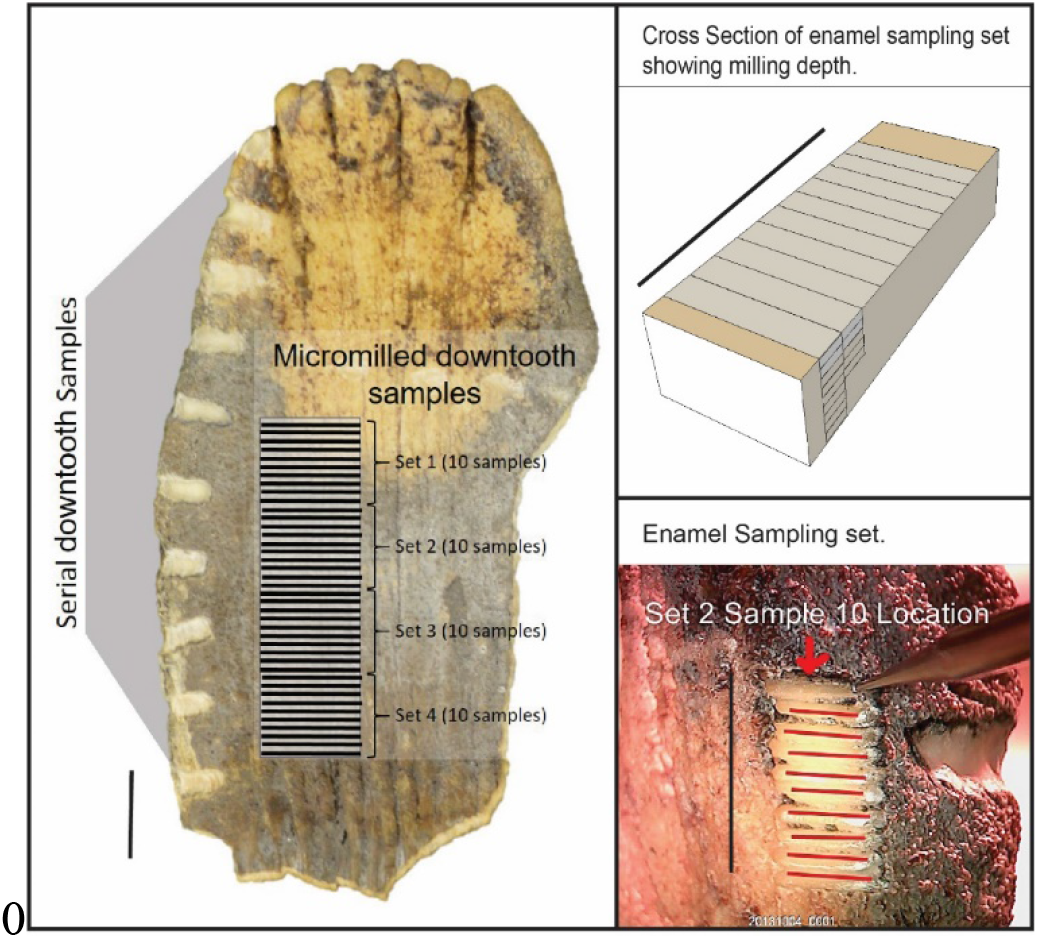
Schematic illustration of serial and micro-sampling strategies. Black bar is equal to 1 cm.

All enamel powder samples were measured in a Finnigan Delta Plus XL mass spectrometer in continuous flow mode connected to a Gas Bench with a CombiPAL autosampler at the Iowa State University Stable Isotope lab, Department of Geological and Atmospheric Sciences. Reference standards (NBS-18, NBS-19) were used for isotopic corrections, and to assign the data to the appropriate isotopic scale. Corrections were done using a regression method using NBS-18 and NBS-19. Isotope results are reported in per mil (‰). The long-term precision (at 1σ) of the mass spectrometer is ±0.09‰ for δ^18^O and ±0.06‰ for δ^13^C, respectively, and precision is not compromised with small carbonate samples (~150 micrograms). Both δ^13^C and δ^18^O are reported VPDB.

Enamel δ^13^C results are corrected −14.1‰ to approximate the δ^13^C of dietary input (δ^13^C_diet_) (Daniel Bryant and Froelich 1995).

Enamel δ^18^O results were converted from VPDB to SMOW using the equation:

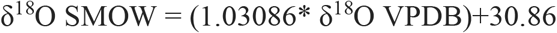

The δ^18^O of enamel phosphate (δ^18^O_p_) for these samples was calculated from the δ^18^O of enamel carbonate (δ^18^O_c_) (Fox and Fisher 2001)

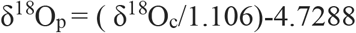

Estimates of body water δ^18^O (δ^18^O_w_) were calculated following Dauxe et al. (2008)

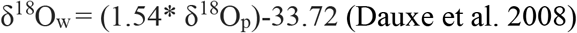

The ^87^Sr/^86^Sr component of enamel bioapatite reflects changes in the geochemical makeup of the surface an animal grazed across during tooth formation. In serial-scale analyses, enamel powder samples were split from the light isotope samples described above, and represent the same portions of mammoth teeth. Each ~5 mg sample of powder was leached in 500 μL 0.1 N acetic acid for four hours to remove diagenetic calcite and rinsed 3 times with deionized water (centrifuging between each rinse). In the micromilled series, small sample sizes prevented Sr analyses from being performed on the same samples as δ^13^C and δ^18^O analyses. Therefore, Sr from these growth series was analyzed opportunistically, or at the scale of 1 sample every 2mm.

All enamel samples were then dissolved in 7.5 N HNO_3_ and the Sr eluted through ion-exchange columns filled with strontium-spec resin at the University of Kansas Isotope Geochemistry Laboratory. ^87^Sr/^86^Sr ratios were measured on a Thermal Ionization Mass Spectrometer (TIMS), an automated VG Sector 54, 8-collector system with a 20-sample turret, at the University of Kansas Isotope Geochemistry laboratory. Isotope ratios were adjusted to correspond to a value of 0.71250 on NBS-987 for ^87^Sr/^86^Sr. We also assumed a value of ^86^Sr/^88^Sr of 0.1194 to correct for fractionation.

The distribution of ^87^Sr/^86^Sr values in vegetation across the surface of the midcontinent is determined by the values of soil parent material. In a large part of this region, surface materials are composed of allochthonous Quaternary deposits such as loess, alluvium, and glacial debris. Therefore continent-scale Sr isoscape models derived from bedrock or water (Bataille and Bowen 2012) are not ideal for understanding first order variability in midwestern ^87^Sr/^86^Sr. For these reasons, Widga et al. (2017) proposed a Sr isoscape for the Midwest based on surface vegetation. At a regional scale, these trends in vegetation ^87^Sr/^86^Sr reflect the Quaternary history of the region, and are consistent with other, empirically derived, patterns in Sr isotope distribution from the area (Slater et al. 2014; Hedman et al. 2018). Sr isotope values in mammoth enamel are compared to a ^87^Sr/^86^Sr isoscape constructed from the combined datasets of Widga et al. (2017b) and Hedman et al. (2009), and Hedman et al. (2018).

## RESULTS

The collagen of 54 mastodons and 22 mammoths was analyzed for δ^13^C_coll_ and δ^15^N_coll_ (Table 2). Three mastodons have ^14^C ages that place them beyond the range of radiocarbon dating, and the youngest mastodons date to the early part of the Younger Dryas, shortly before extinction. Despite being well-represented prior to the LGM and during deglaciation (i.e., Oldest Dryas, Bølling, Allerød, Younger Dryas), this dataset lacks mastodons from the study region during the coldest parts of the LGM. Mammoths are present in this dataset from 40 ka until their extinction in the region during the late Allerød.

Visually, there is substantial amounts of overlap in the δ^13^C_coll_ and δ^15^N_coll_ of mammoths and mastodons through the duration of the dataset (Figure 3, Table 2). Both taxa show average δ^13^C_coll_ values around −20‰, consistent with a diet dominated by C3 trees, shrubs, and/or cool-season grasses. This is broadly consistent with variable, but shared diets during time periods when both taxa occupied the region. The average δ^13^C_coll_ values for mammoths (−20.4‰) is similar to that of mastodons (−21.0‰). The average δ^15^N_coll_ of mammoths (7.5‰) is elevated compared to mastodons (4.4‰).

**Figure 3.**
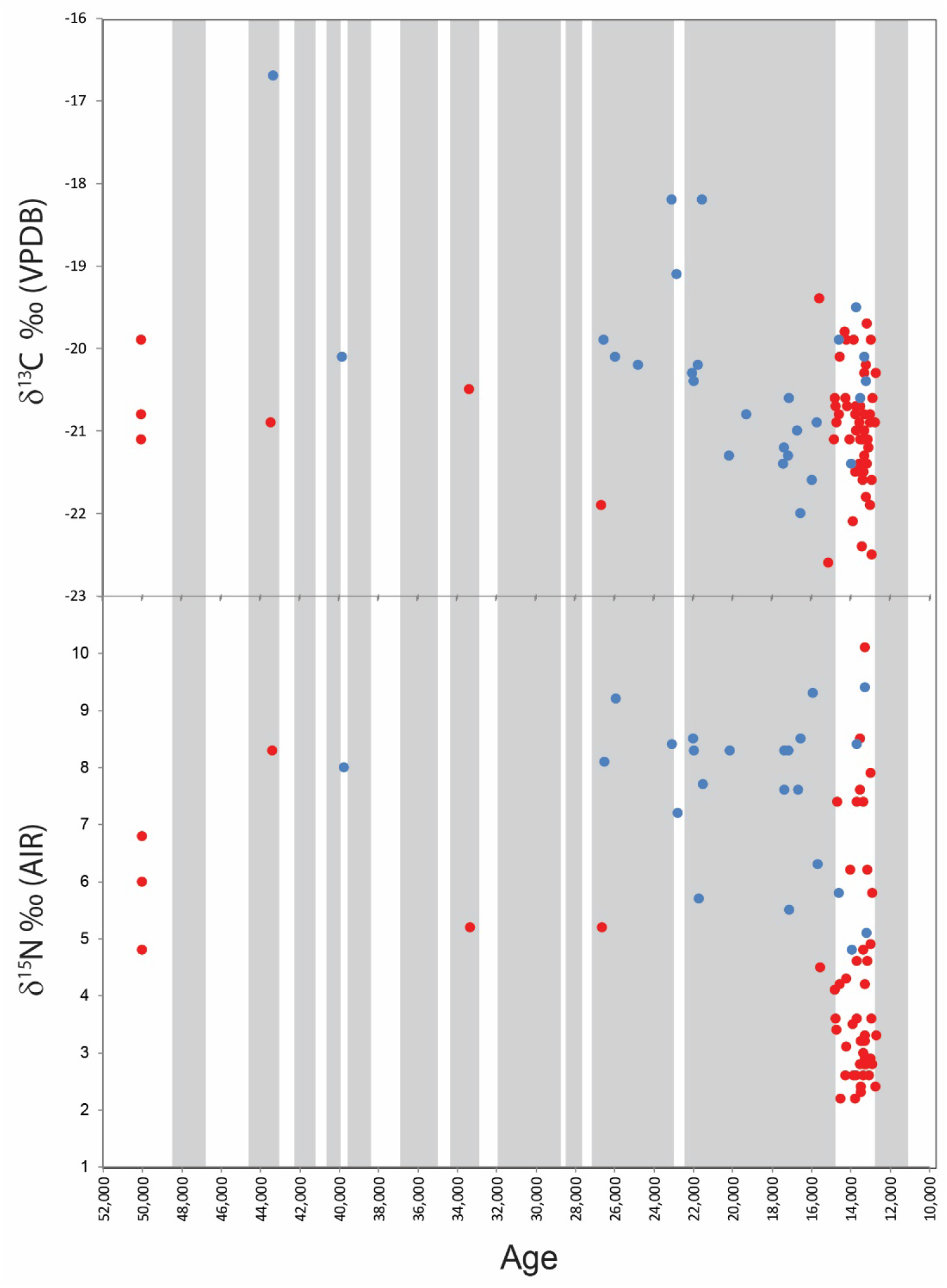
Changes in midwestern proboscidean collagen δ^13^C and δ^15^N through time. Mastodons=red, Mammoths=blue, glacial stadials = grey bars. Stadial chronology from Rasmussen et al., 2014.

However, dietary relationships between taxa are not static through time. Prior to the LGM, both δ^13^C_coll_ and δ^15^N_coll_ are significantly different between mammoths and mastodons (δ^13^C t-test; p=0.013; δ^15^N t-test; p=0.010). During the Oldest Dryas the δ^13^C_coll_ of both taxa is very similar, although δ^15^N_coll_ between mammoths and mastodons remains distinct (t-test; p=0.012). During the Allerød, δ^13^C_coll_ and δ^15^N_coll_ between taxa are indistinguishable.

The δ^13^C_coll_ signature of mastodon diets changes little throughout the last 50 ka. Despite a noticeable absence of mastodon material during the height of the LGM, the only significant shift in the δ^13^C_coll_ of mastodon diets occurs between the Bølling and the Allerød (t-test; p=0.008).

Mastodon δ^15^N_coll_ values fall clearly into two groups, those that date prior to the LGM, and those that post-date the LGM. Mastodon δ^15^N_coll_ values are significantly higher in pre-LGM samples than in samples dating to the Younger Dryas (t-test; p=0.010), Allerød (t-test; p=0.017), and Bølling (t-test; p<0.000). Of note is a group of mastodons that show lower δ^15^N_coll_ values during the Oldest Dryas, Bølling, Allerød, and Younger Dryas, when the average δ^15^N_coll_ values decrease to values <5‰. A similar shift is not evident in mammoths at this time.

Mammoth δ^13^C_coll_ during MIS 3 is significantly different from mammoths dating to LGM II (t-test; p=0.008) or the Oldest Dryas (t-test; p=0.005). There are no significant differences in mammoth δ^15^N_coll_ between different time periods.

All serial enamel series were between 9 and 16 cm in length and represent multiple years. Stable oxygen isotopes of three MIS 2 mammoths (Principia College, Brookings, Schaeffer) overlap significantly, while MIS 4 (234JS75) and MIS 5e (232JS77) mammoths from Jones Spring show elevated values indicative of warmer conditions (Table 3; Figure 4). The variance in serially sampled MIS 2 mammoths (average amp. = 2.5‰) was not significantly different from the series from Jones Spring, dating to MIS 4 (amp. = 2.9‰)(F-test; p=0.320). However, the MIS 5e mammoth from Jones Spring has a smaller amplitude (amp. = 1.1‰) compared to both the MIS 4 mammoth (F-test; p=0.003) and the three MIS 2 mammoths (F-test; p<0.000). Strontium isotope ratios of this animal indicates a home range on older surfaces of the Ozarks, throughout tooth formation, with a period of relatively elevated ^87^Sr/^86^Sr values also corresponding to low δ^18^O values.

**Figure 4.**
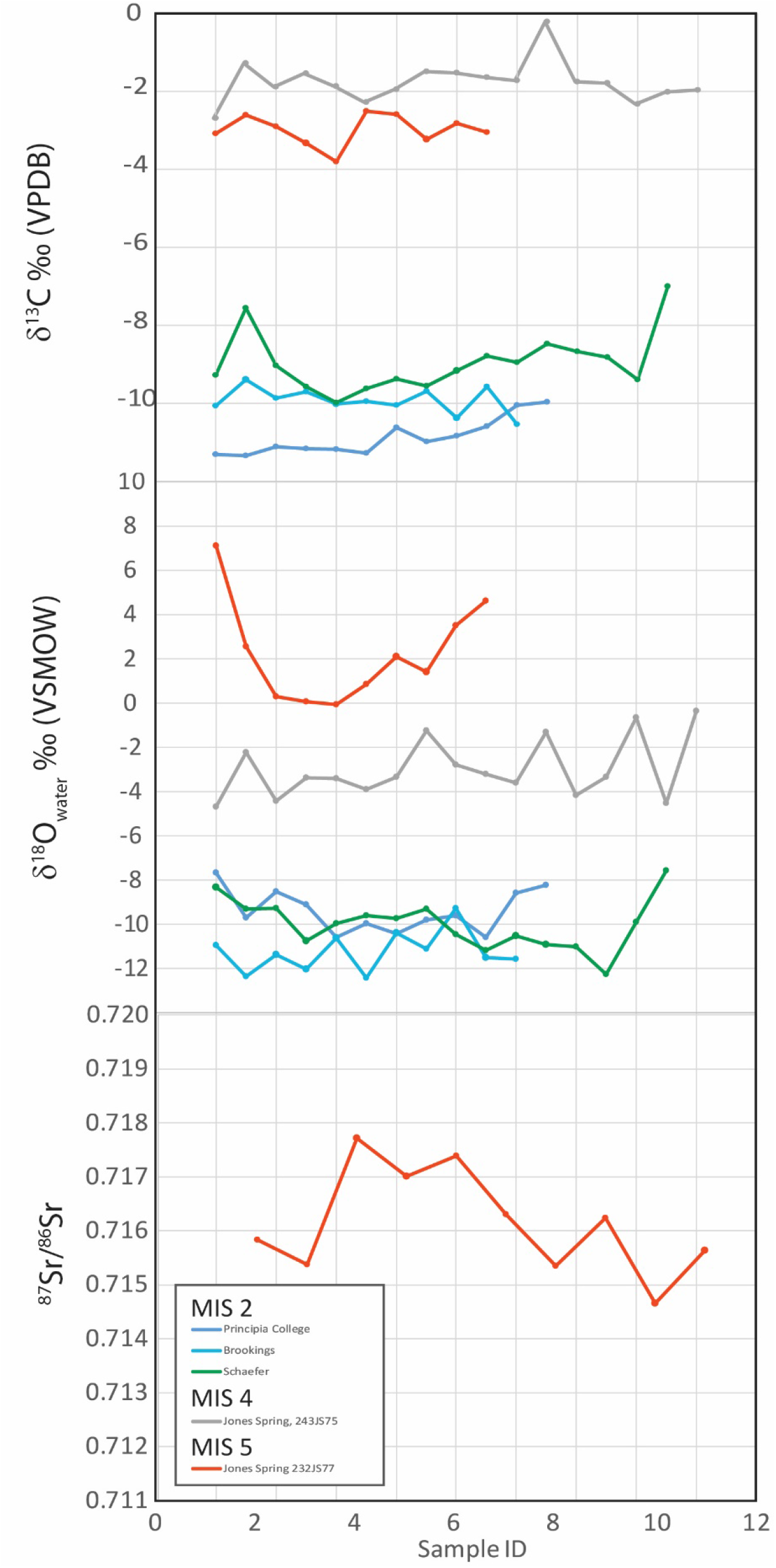
Time series, serial enamel δ^13^C, δ^18^O, ^87^Sr/^86^Sr.

Two mammoth teeth from the Jones Spring locality in Hickory Co., MO were microsampled (Figure 5). These specimens are from beds representative of MIS 5e and MIS 3 deposits.

**Figure 5.**
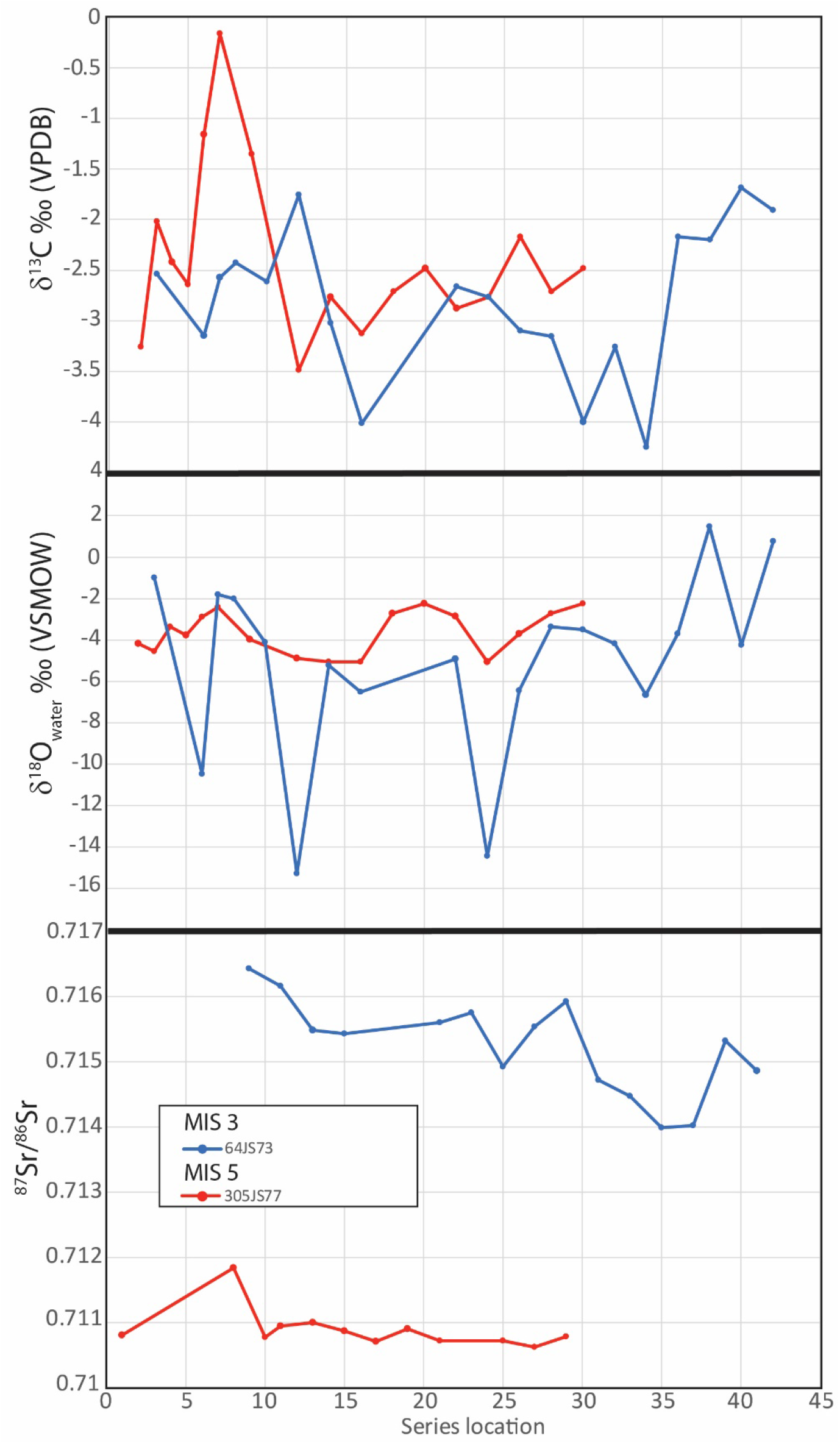
Time series, micro-sampled enamel δ^13^C, δ^18^O, ^87^Sr/^86^Sr. Both specimens are from Jones Spring, Hickory County, Missouri.

The MIS 3 molar (64JS73) shows regular negative excursions in δ^18^O values suggestive of seasonal temperature changes in ingested water. These excursions are relatively short-lived and extreme. Seasonal variation in enamel growth rate and maturation may account for the perceived short length of these periods. Maximum δ^18^O values in both mammoths are similar however; the MIS 5e mammoth lacks these negative excursions and exhibits values that are more complacent through the length of the tooth.

The δ^13^C series from both molars indicate these mammoths experienced a C4 diet throughout the year. However, deep dips in the δ^18^O series of the MIS 3 molar also correspond to at least two temporary peaks in the δ^13^C series.

The ^87^Sr/^86^Sr isotope series from both animals demonstrate an adherence to separate home ranges. The MIS 5e mammoth shows values similar to bedrock units outcropping locally in central-western Missouri and neighboring areas of Kansas and Oklahoma. The MIS 3 mammoth, however, had a home range across rock units with much higher ^87^Sr/^86^Sr values. The home ranges for these animals do not overlap, despite their recovery from different strata within the same locality.

## DISCUSSION

Despite overall similar isotopic values in mammoths and mastodons throughout the period of this study, underlying nuances are informative to regional changes in niche structure and climate. The absence of mastodons during the coldest parts of the LGM suggests that *Mammut* were, at least to some degree, sensitive to colder climates and adjusted its range accordingly.

Comparisons between co-eval mammoths and mastodons indicate a gradual collapse of niche structure. Prior to MIS 2, δ^13^C_coll_ and δ^15^N_coll_ values between taxa were significantly different. By the end of the Allerød, mammoth and mastodon diets were isotopically indistinguishable.

Through time, there was no significant change in the δ^13^C_coll_ of mastodons. Although average mammoth δ^13^C_coll_ in the region during the latter part of the LGM was slightly more negative than other time periods, this is not a marked shift and may be a function of decreased pCO2 during that time (Schubert and Jahren 2015). Globally, Siberian and European mammoths exhibit a similar range of δ^13^C_coll_ values (Iacumin et al. 2010; Szpak et al. 2010; Arppe et al. 2019; Schwartz-Narbonne et al. 2019).

In both taxa, mean δ^15^N_coll_ decreases slightly throughout the sequence, however, the minimum δ^15^N values for mastodons during the Bølling, Allerød, and Younger Dryas are significantly lower than earlier mastodons. The anomalously low values of these late mastodons are also shared with other published mastodon values in the Great Lakes region (Metcalfe et al. 2013). They are also significantly lower than contemporary midwestern, Eurasian or Beringean mammoths (with some exceptions, Drucker et al. 2018). The timing of these low, mastodon δ^15^N_coll_ values correspond to regionally low δ^15^N_coll_ values in the bone collagen of non-proboscidean taxa from European late Quaternary contexts (Drucker et al., 2009; Richards and Hedges, 2003; Rabanus-Wallace et al., 2017; Stevens et al., 2008) suggesting broad changes in global climate may have had cascading impacts in the N budget of local ecosystems.

However, understanding in more detail how climate might have affected N cycling in midwestern ecosystems remains unclear. Flux in soil and plant N may be a function of plantbased N2 fixation (Shearer and Kohl 1993), rooting depth (Schulze et al. 1994), N loss related to climate factors (Austin and Vitousek 1998; Handley and Raven 1992), microbial activity, or mycorrhizal colonization (Hobbie et al. 2000; Michelsen et al. 1998;). Furthermore, the role of large herbivore populations in N flux may be significant (Frank et al. 2004). In N limited environments such as tundra, terrestrial plants may receive relatively more N from inorganic sources. Boreal forests like those of the Midwest during the Bølling and Allerød, however, exhibit relatively greater biological productivity, so plant shoots are more likely to take in volatized ammonia (low δ^15^N) from organic sources such as urea (Fujiyoshi et al. 2017).

At this time, it is difficult to distinguish which (if any) of these factors had an impact on late Quaternary mastodon δ^15^N_coll_ values. Although there is a wide range in δ^15^N_coll_ of mammoths and mastodons throughout the sequence, δ^15^N_coll_ values below 5‰ are only present in mastodons during post-LGM deglaciation in the lower Great Lakes (39-43 deg. latitude), throughout an area suggested to be vegetated by a disharmonius flora dominated by Black Ash and Spruce (Gonzales and Grimm 2009; Gill et al. 2009) (Figure 7). The low δ^15^N_coll_ values suggest that these mastodons occupied an undefined, local-scale, dietary niche that was not shared by contemporary mammoths, or by earlier mastodons. Previous research has suggested that the Bølling-Allerød may have been a time of high mastodon populations in the Great Lakes region (Widga et al. 2017a). The concentration of mastodons around water sources may have had an impact on the δ^15^N of browsed plants due to increased contributions of herbivore urea. Isotopically light plant shoots can occur with increased utilization of volatized NH4 from organic sources, including urea from herbivores.

**Figure 7.**
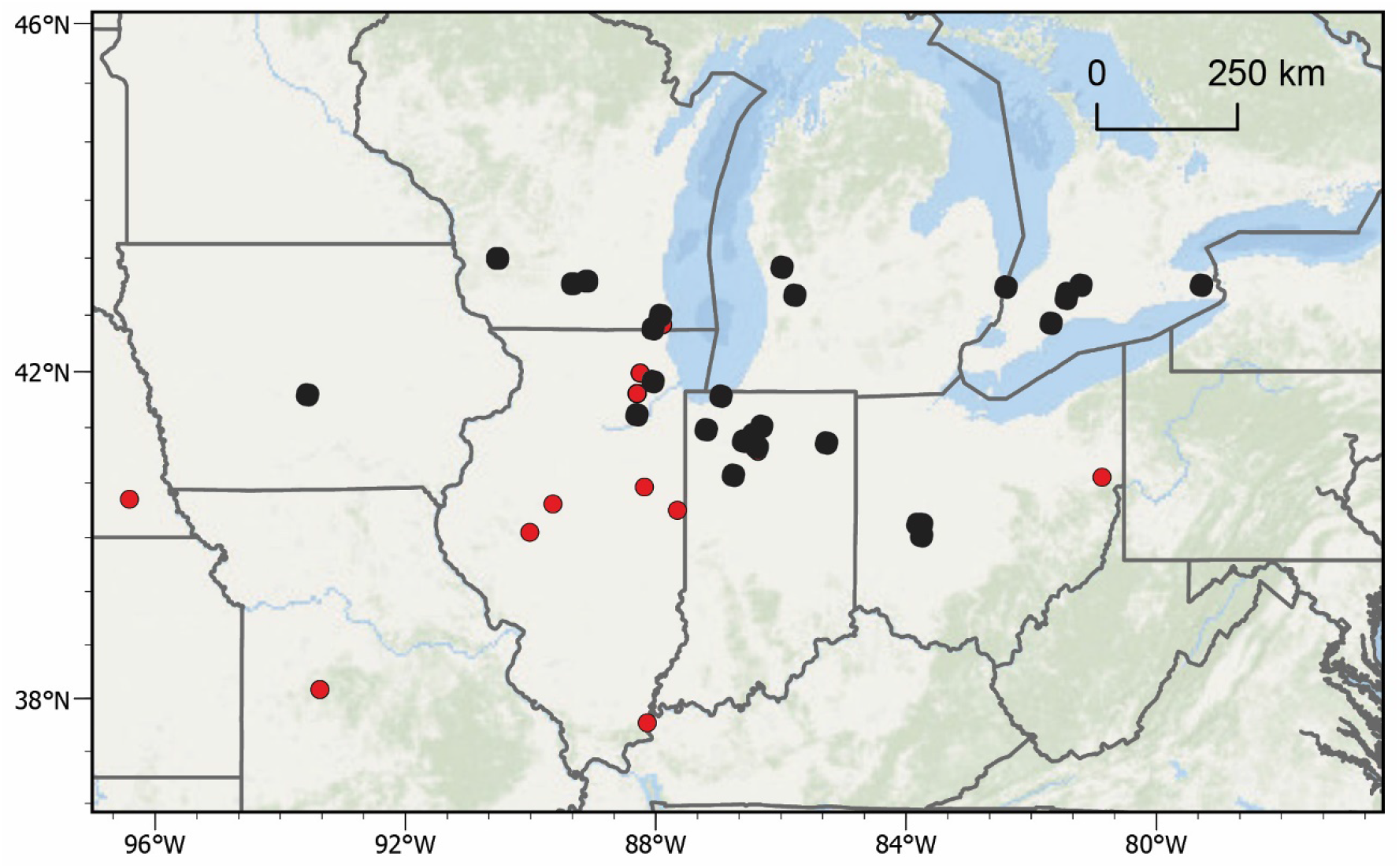
Mastodon distribution during the Bølling-Allerød. Black circles indicate mastodon δ^15^N_coll_ < 5‰. Red circles represent mastodons with δ^15^N_coll_ > 5‰.

### Life Histories of midwestern Mammoths

Through analyses of incremental growth structures in mammoths, such as molar enamel or tusk dentin, we can reconstruct longitudinal life histories that reflect the landscape experienced by an animal over multiple years. Late Pleistocene mammoths from the study area exhibit relatively consistent, down-tooth patterns in δ^13^C and δ^18^O. Similar bulk enamel isotope values in South Dakota, Wisconsin, and Illinois mammoths suggest access to broadly similar resources, and relatively stable access to these resources across multiple years of the life of an individual (Figure 4). Less negative δ^13^C and δ^18^O values in the last forming samples of the Schaeffer mammoth suggest a change in that animal’s life history in the year before death. This change could be explained by local environmental changes resulting in nutritional stress.

Two mammoths from the Jones Spring site in southwest Missouri provide a pre-LGM perspective on landscapes that mammoths occupied (Figure 4). A tooth plate dating to MIS 5e from Jones Spring has less negative δ^18^O and δ^13^C values compared to the MIS 2 samples. This indicates warmer overall conditions and a diet that incorporated more C4 grasses throughout multiple years. Pollen from the same stratigraphic units further suggest that southwestern Missouri during MIS 5e was dominated by *Pinus* (no *Picea)* with a significant non-arboreal component (King 1973). Importantly, Sr isotopes from this tooth indicate that while this tooth was forming, the animal was foraging across surfaces that are more radiogenic than local values. The nearest area with ^87^Sr/^86^Sr values >0.7140 is the central Ozark uplift to the east of Jones Spring (Figure 6). The wide amplitude of δ^18^O values throughout the length of this tooth, combined with the Sr isotope data suggesting adherence to the central Ozark uplift suggest relatively broad shifts in annual water availability and/or that this animal utilized a variety of water sources, including surface sources and freshwater springs.

**Figure 6.**
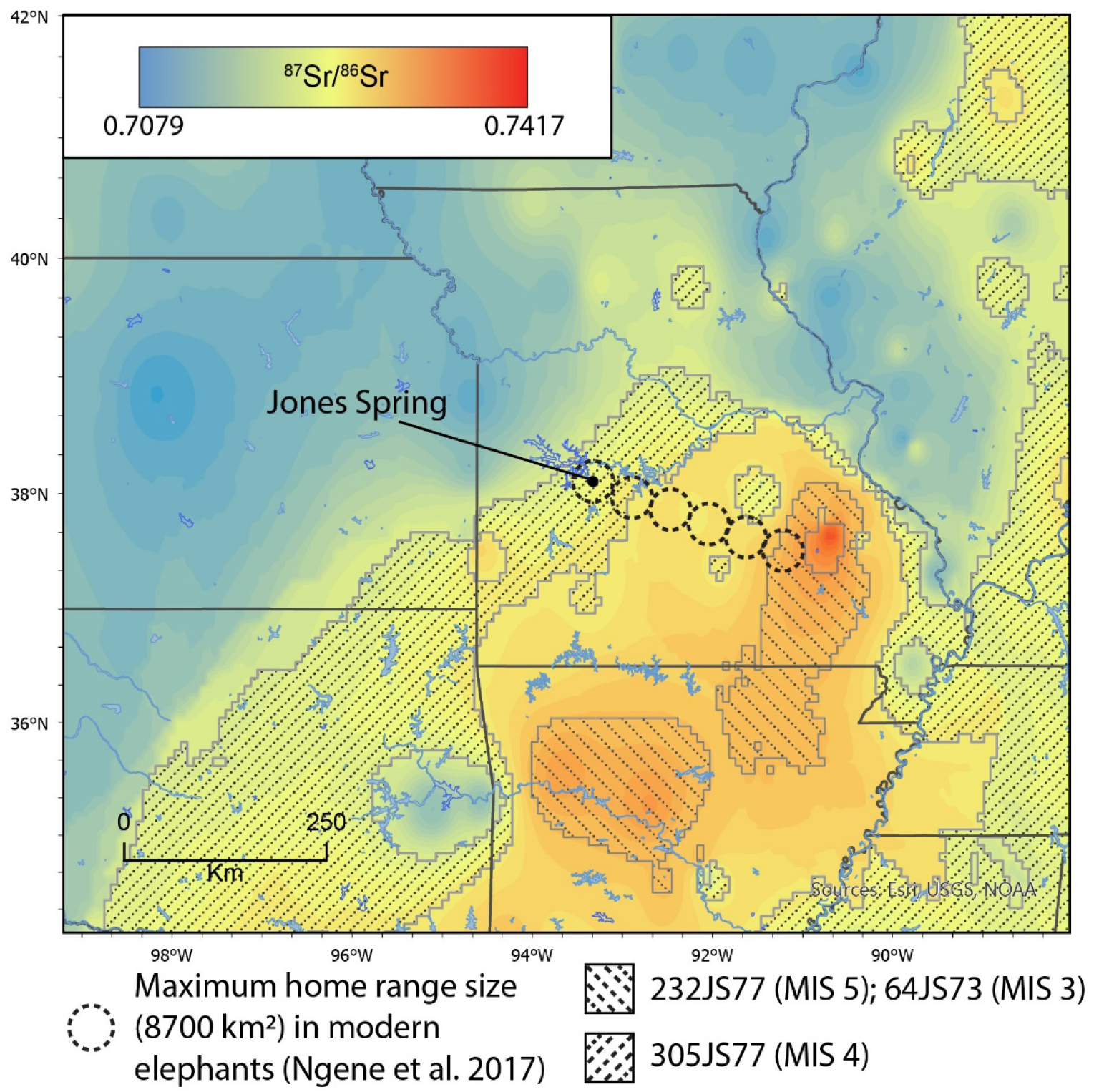
Mobility in mammoths from Jones Spring, Hickory Co., MO. Hatched areas indicate the range of ^87^Sr/^86^Sr values from mammoth molar ridge-plates. One mammoth (305JS77) exhibits local values. Two separate mammoths (232JS77, 64JS73) exhibit Sr values suggesting >200 km movement from the central core of the Ozark uplift. This implies movement over multiple years that is at least six times larger than the maximum home range size documented in modern elephants (Netosha National Park, Namibia). Basemap isoscape data from Widga et al. (2017b), Hedman et al. (2009, 2018).

The MIS 4 molar from Jones Spring has δ^18^O values intermediate between the MIS 2 samples and the MIS 5e sample, along with δ^13^C values that indicate more C4 consumption than MIS 2 samples (Figure 4). This mammoth occupied a cooler environment than the MIS 5e mammoth, but significantly warmer than the late Pleistocene mammoths in the MIS 2 group.

In the case of the two, microsampled mammoths from Jones Spring, MO, isotopic trends in each animal illustrates different life histories (Figure 5). The MIS 4 mammoth occupied the western Ozarks as indicated by ^87^Sr/^86^Sr values deposited throughout the development of the sampled portion of the tooth. It occupied a landscape where C4 vegetation was common, with mild winter temperatures. The MIS 3 mammoth from Jones Spring occupied the central core of the Ozark uplift indicated by ^87^Sr/^86^Sr values deposited during the formation of the sampled tooth. However, between the cessation of enamel formation and death, this animal moved ~200 km to the western Ozarks where it was preserved in the Jones Spring deposits. The δ^18^O series of this animal suggests greater seasonality during MIS 3 compared to MIS 5e, with deep negative excursions during the cold season. It also had a diet composed primarily of C4 vegetation.

These results suggest a broadly similar geographic scale of landscape-use in late Quaternary mammoths in the Midwest to mammoths in the Great Plains and Florida (Esker et al. 2019; Hoppe et al. 1999; Hoppe 2004). Overall, mammoths from both regions do not engage in significant seasonal migrations, but can move greater distances at annual to decadal time-scales. Further, although δ^13^C_coll_ values in mammoths clearly suggest a niche that, at times, included C3-rich diets in a region that was dominated by forest, not grasslands, both Jones Spring mammoths were mixed feeders with C4 grasses making up a significant part of their diet. The individual life histories of these mammoths are highly variable, underscoring the need to control for individual mobility in stable isotope studies of large herbivore taxa.

### Testing scenarios of late Pleistocene Proboscidean population dynamics

Can this stable isotope dataset illuminate trends in late Proboscidean population dynamics during deglaciation in the Midwest? It is possible that the shift in δ^15^N represents the colonization of a novel ecological niche that also coincides with climate and vegetational changes at the beginning of the Bølling-Allerød. However, it is unclear what this niche might be. Stable isotopes alone do not adequately define this niche space, and further paleobotanical work is necessary. Drucker et al. (2018) also noted anomalously low δ^15^N values in LGM mammoths from Mezyhrich in central Europe, and attributed this pattern to a large, but unspecific, shift in dietary niche. This is counter-intuitive in warming landscapes of the Midwest since boreal forests typically have increased amounts of N fixing microbes relative to tundra environments, thus increasing the δ^15^N in forage, overall.

Some studies have suggested that the growth rate of an individual is inversely correlated with δ^15^N (Warinner and Tuross 2010). This would be consistent with some scenarios of late Pleistocene mastodon population dynamics. Fisher (Fisher 2009; 2018) suggests that predator pressure from Paleoindian groups who were megafaunal specialists would have caused mastodons to mature at a younger age. A decrease in the age at weaning would mean a shorter period of nursing-related, elevated dietary δ^15^N in young mastodons. If this were the case, we would expect an overall decrease in δ^15^N of bone collagen in animals in their first and second decade of life. In our dataset, there is no significant change in maximum or mean δ^15^N in mastodon bone collagen, despite a subset of mastodons that have lower δ^15^N values. If predator pressure is contributing to faster maturation and shortening the time of nursing, then it is only occurring in some areas. However, even if this were the case in these areas, it is still uncertain what ecological processes might drive an increase in growth rate. Depending on local forage conditions experienced by an animal, an increase in growth rate may be caused by increased predator pressure (Fisher 2009) or a decrease in population density (Wolverton et al. 2009).

It is also possible that low δ^15^N values in post-LGM mastodons represents a systematic change in predator avoidance strategies among some mastodon populations (Fiedel et al. 2019). Mastodons may have been attracted to low δ^15^N areas such as marshes and wetlands as a response to new predators on the landscape (i.e., humans). However, taphonomic modification of late Pleistocene proboscidean materials by humans or other predators is extremely rare in this dataset (Widga et al. 2017a), so we see this scenario as an unlikely driver of mastodon landscape use.

If proboscidean populations are dense on the landscape (Widga et al. 2017a) there may also be rapid, significant, and long-term changes to N cycle. An influx of N via urea as would be expected in high-density areas, would cause isotopically heavy plant roots and isotopically light shoots. Under intense grazing pressure, changes to N contributions change quickly from inorganic soil mineral N reservoirs to N from urea (McNaughton et al. 1988; Knapp et al. 1999). Individuals with lower δ^15^N_coll_ values during the Allerød lived in areas with high mastodon populations, which may have contributed to isotopically light forage (due to ammonia volitization, Knapp et al. 1999). However, this scenario does not explain why this shift is absent in mammoths during this time.

## CONCLUSIONS

Stable isotopes of proboscidean tissues in the midwestern US illustrate the variability inherent in modern paleoecological analyses. Traditional approaches to understanding animal diets and landscape use through morphology and a reliance on modern analogues is inadequate for understanding paleoecological changes within megafaunal populations, or during times of rapid ecological change such as the late Quaternary.

Stable isotopes from bone collagen of midwestern proboscideans suggest consistently C3-dominated diets over the last 50 ka. During the LGM, mammoth diets may have included C3 grasses, but the prevalence of C3 flora during the post-LGM period is likely due to a landscape shift to more forest (Gonzales and Grimm 2009; Saunders et al. 2010) with grasses making up very little of the floral communities in the southern Great Lakes region.

Despite the strong C3-signal in mammoth and mastodon diets during this period, the range of dietary flexibility and the degree of overlap between these two taxa is striking. The isotopically defined dietary niche of mammoths and mastodons show increasing overlap as they approach extinction. Niche overlap is also supported by assemblages where both taxa co-occur (Saunders et al. 2010; Widga et al. 2017a). However, there are exceptions to this pattern and some late mastodons exhibit low δ^15^N values that may indicate the evolution and occupation of a new dietary niche, physiological responses to late Pleistocene ecological changes, or some other process acting on late mastodon populations.

Further resolution of mammoth and mastodon life histories are gleaned from stable isotopes in tooth enamel. δ^13^C and δ^18^O generally track climate and landscape changes experienced during tooth formation. MIS 2 mammoths are broadly similar in their isotopic life histories and illustrate relative homogeneity of landscape conditions across the Midwest at this time. Micro-sampled mammoth molars from Jones Spring, MO, indicate a significant increase in MIS 3 seasonality, compared to the last interglacial period (MIS 5e). Finally, some mammoths in this study died >200 km away from where they lived during the formation of sampled molars suggesting lifetime mobility patterns in mammoths could have a significant effect on presumed ‘local’ stable isotope values.

## Acknowledgements

The study benefited from discussions with Paul Countryman, Eric Grimm, and Bonnie Styles, then at the Illinois State Museum; Rhiannon Stevens (University College of London) and Matt G. Hill (Iowa State University). Additional conversations with Don Esker greatly clarified micro-milling methods, and Joseph Andrew (University of Kansas) assisted with Sr isotope analyses. Jeff Pigati and two anonymous reviewers provided invaluable feedback for strengthening the manuscript. This research was funded by NSF grants 1050638, 1049885 and 1050261 and the Illinois State Museum Society. Finally, we would like to acknowledge all of the collections managers and curators who assisted us with access to the specimens within their care.

## Tables

Table 1. Temporal Scale in Paleoecological Analyses.

Table 2. Mammoth and mastodon collagen stable isotope values, by chronzone.

Table 3. Summary of serial and micro-sampled tooth enamel series: δ^13^C, δ^18^O, ^87^Sr/^86^Sr.

## Supplemental information

SM Supplemental Material. Word File

SM Table 1. Mammoth and Mastodon δ^13^C, δ^15^N data.

SM Table 2. Mammoth enamel isotope data.

SM Table 3. Radiocarbon dated mammoths and mastodons from the Midwest.

