## Supplementary material for "Life histories and niche dynamics in late Quaternary proboscideans from Midwestern North America: evidence from stable isotope analyses": SM

Supplemental Materials

*Data quality*. We undertook a series of analyses to ensure that our methods were appropriate for the questions that were being asked.

1. Impact of diagenesis on micromilled samples. Sample loss was a major concern due to the small size (~1mg) of micromilled enamel samples. For this reason, we chose to forego traditional pretreatment of these samples where there was the possibility of sample loss (or reduction) due to handling. Instead, we analyzed paired treated/untreated bulk enamel samples to assess the impact of diagenesis on isotopic results (SM Table 2). Untreated bulk enamel samples were consistently <1‰ different from enamel samples treated with 0.1N acetic acid and 2.5% NaOCl. Therefore we expect that diagenetic alteration of enamel carbonate to have little impact on δ13C, δ18O, and 87Sr/86Sr results.

SM Table 2. Results of paired, NaOCl+Acetic and Water only samples.

|  | NaOCl+Acetic | | Water only | | Difference | |
| --- | --- | --- | --- | --- | --- | --- |
| Sample | δ13C (VPDB) | δ18O (VSMOW) | δ13C (VPDB) | δ18O (VSMOW) | d13C | d18O |
| 64JS73, sample 3 | -0.25 | 27.65 | -0.79 | 27.58 | 0.54 | 0.07 |
| 64JS73, sample 8 | -3.45 | 23.56 | -3.71 | 23.52 | 0.26 | 0.04 |

2. Impact of sample depth on micromilled samples. The timing of enamel growth and maturation complicates analyses of sub-annual isotopic changes across many species of large herbivores ^1,2^. This has been demonstrated to be an issue for mammoth enamel ridge-plates as well ^3^. To assess the effectiveness of our micromilling method, we analyzed three depth profiles for δ13C and δ18O (SM Table 3). There are significant changes in both isotopes between outer and inner enamel samples (SM Figure 1). There is no evidence for significant diagenetic alteration, so we assume these trends reflect the timing of enamel maturation, with the latest forming enamel forming at inner boundary between enamel and dentin.

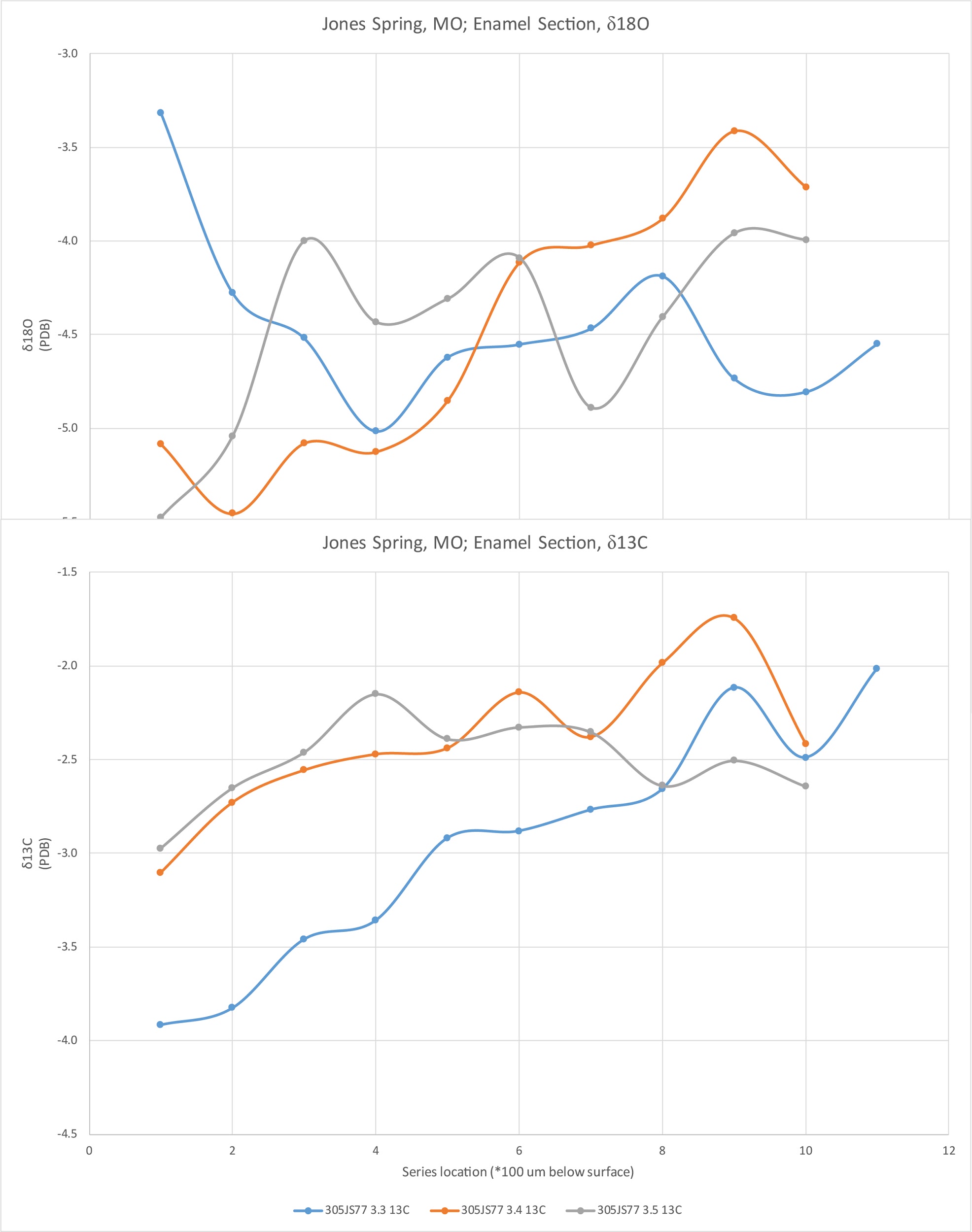

SM Figure 1. Depth profiles of stable isotope values through the thickness of mammoth tooth enamel.
